# Phototrophic lactate utilization by *Rhodopseudomonas palustris* is stimulated by co-utilization with additional substrates

**DOI:** 10.1101/502963

**Authors:** Alekhya Govindaraju, James B. McKinlay, Breah LaSarre

**Affiliations:** Microbiology and Biotechnology undergraduate programs, Indiana University, Bloomington, IN 47401; Department of Biology, Indiana University, Bloomington, IN 47401

**Keywords:** mixed substrate utilization, *Rhodopseudomonas palustris*, lactate, catabolite repression, diauxie

## Abstract

The phototrophic purple nonsulfur bacterium *Rhodopseudomonas palustris* is known for its metabolic versatility and is of interest for various industrial and environmental applications. Despite decades of research on *R. palustris* growth under diverse conditions, patterns of *R. palustris* growth and carbon utilization with mixtures of carbon substrates remain largely unknown. *R. palustris* readily utilizes most short chain organic acids but cannot readily use lactate as a sole carbon source. Here we investigated the influence of mixed-substrate utilization on phototrophic lactate consumption by *R. palustris*. We found that lactate was simultaneously utilized with a variety of other organic acids and glycerol in time frames that were insufficient for *R. palustris* growth on lactate alone. Thus, lactate utilization by *R. palustris* was expedited by its co-utilization with additional substrates. Separately, experiments using carbon pairs that did not contain lactate revealed acetate-mediated inhibition of glycerol utilization in *R. palustris*. This inhibition was specific to the acetate-glycerol pair, as *R. palustris* simultaneously utilized acetate or glycerol when either was paired with succinate or lactate. Overall, our results demonstrate that (i) *R. palustris* commonly employs simultaneous mixed-substrate utilization, (ii) mixed-substrate utilization expands the spectrum of readily utilized organic acids in this species, and (iii) *R. palustris* has the capacity to exert carbon catabolite control in a substrate-specific manner.

**IMPORTANCE:** Bacterial carbon source utilization is frequently assessed using cultures provided single carbon sources. However, the utilization of carbon mixtures by bacteria (i.e., mixed-substrate utilization) is of both fundamental and practical importance; it is central to bacterial physiology and ecology, and it influences the utility of bacteria as biotechnology. Here we investigated mixed-substrate utilization by the model organism *Rhodopseudomonas palustris*. Using mixtures of organic acids and glycerol, we show that *R. palustris* exhibits an expanded range of usable carbon substrates when provided in mixtures. Specifically, co-utilization enabled the prompt consumption of lactate, a substrate that is otherwise not readily used by *R. palustris*. Additionally, we found that *R. palustris* utilizes acetate and glycerol sequentially, revealing that this species has the capacity to use some substrates in a preferential order. These results provide insights into *R. palustris* physiology that will aid the use of *R. palustris* for industrial and commercial applications.

## INTRODUCTION

Many bacteria in natural environments likely consume multiple carbon sources simultaneously (1, 2). However, bacterial substrate utilization is most often studied using bacteria in isolation with single carbon sources (3). Data on mixed-substrate utilization in diverse bacteria is crucial for both the understanding of nutrient acquisition, metabolism, and community dynamics within microbial ecosystems (1, 4) and the rational application of bacteria as biotechnology (5-8).

When encountering multiple carbon sources (i.e., substrates), a bacterium will utilize the substrates either simultaneously (i.e., co-utilization) or sequentially depending on the identity of the substrates. Sequential utilization typically results in a diauxic growth pattern characterized by two or more exponential phases that are each separated by a lag phase (2, 9); however, sequential utilization can also occur without an intervening lag phase (2, 4, 10), a pattern often referred to as “biphasic growth.” It generally holds true that, during sequential utilization, bacteria preferentially use the carbon source that supports the highest growth rate during the first phase of growth while utilization of the other carbon source(s) is limited until the preferred carbon source is no longer available (2); this process is commonly referred to as carbon catabolite repression (CCR) (11). A classic example of CCR involves the *lac* operon in *Escherichia coli*. This operon, which is required for lactose transport and metabolism, is only transcribed when lactose is present and glucose is absent, resulting in the preferential consumption of glucose prior to lactose when cells are provided a mix of the two sugars (11). Although sequential carbon utilization was once thought to predominate among bacteria, particularly at high substrate concentrations (2-4), it has become clear that simultaneous utilization of carbon substrates is common (1). For example, co-utilization occurs in *Pseudomonas putida* grown with glucose and aromatic compounds (12), in *E. coli* grown with various organic acid pairs (13), and in *Lactococcus brevis* grown with glucose paired with other sugars (14).

The phototrophic purple nonsulfur bacterium *Rhodopseudomonas palustris* is a model organism for investigating metabolic flexibility in response to environmental conditions (15, 16) and is of interest for various commercial applications, including the production of hydrogen gas (17-19) and the biodegradation of aromatic inhibitors in biofuel feedstocks (20). However, there is limited information regarding carbon preference, mixed-substrate utilization, or carbon catabolite control in this species. We recently showed that *R. palustris* utilizes multiple products of *E. coli* mixed-acid fermentation when grown in a synthetic coculture (21). One of these products, lactate, is not readily utilized as a sole carbon source by phototrophically grown *R. palustris* (18). Taken together, these observations raised the question, how does coculturing enable *R. palustris* lactate consumption? Using mixtures of different carbon substrates, here we show that *R. palustris* simultaneously consumes lactate with several other organic acids and glycerol. Importantly, this co-utilization endowed *R. palustris* with the ability to promptly utilize lactate, despite that lactate alone did not support growth in the same time frame. Hence, in coculture, the presence of mixed-acid fermentation products enables the co-utilization of lactate by *R. palustris*. Separately, experiments with additional carbon pairings revealed acetate-mediated inhibition of glycerol catabolism, establishing that *R. palustris* can exert hierarchical regulation of carbon utilization.

## RESULTS

### Acetate and succinate prompt the expedited and simultaneous utilization of lactate

We previously observed that, when grown anaerobically in a mutualistic coculture with *E. coli*, *R. palustris* simultaneously consumed the acetate, succinate, and lactate excreted as fermentation products by *E. coli* during the 200 h culturing period (21). Whereas acetate and succinate are both known to be readily utilized by *R. palustris* (19), *R. palustris* requires long-term incubation with lactate before it will utilize lactate as a sole carbon source (18). Indeed, we observed no growth of *R. palustris* with lactate alone within the time frames sufficient for growth on other organic acids (≤ 300 h; see below). As the utilization of a given carbon source can be influenced by the presence of an additional carbon source (1, 12, 13), we hypothesized that the presence of mixed fermentation products facilitated lactate utilization by *R. palustris* in coculture. We reasoned that the most likely mediators of this effect would be succinate and/or acetate given that they were the other carbon substrates consumed by *R. palustris* in coculture. To test if the presence of acetate and succinate could stimulate lactate consumption, we grew *R. palustris* Nx, the strain we had used in coculture with *E. coli*, in monoculture with a mixture of succinate, acetate, and lactate (5 mM each). Cultures grown with this mix exhibited a single exponential phase with a specific growth rate of 0.074 ± 0.001 h^-1^ (± SD), and growth plateaued within 150 h (**Fig. 1A**). To determine which substrate(s) in the mix had been consumed, we analyzed culture supernatants using high-performance liquid chromatography (HPLC). The data showed that all three compounds had been partially consumed by mid-log-phase and were fully consumed by stationary phase (**Fig. 1B**). Thus, acetate and succinate were sufficient to expedite lactate consumption by *R. palustris*.

**Fig. 1.**
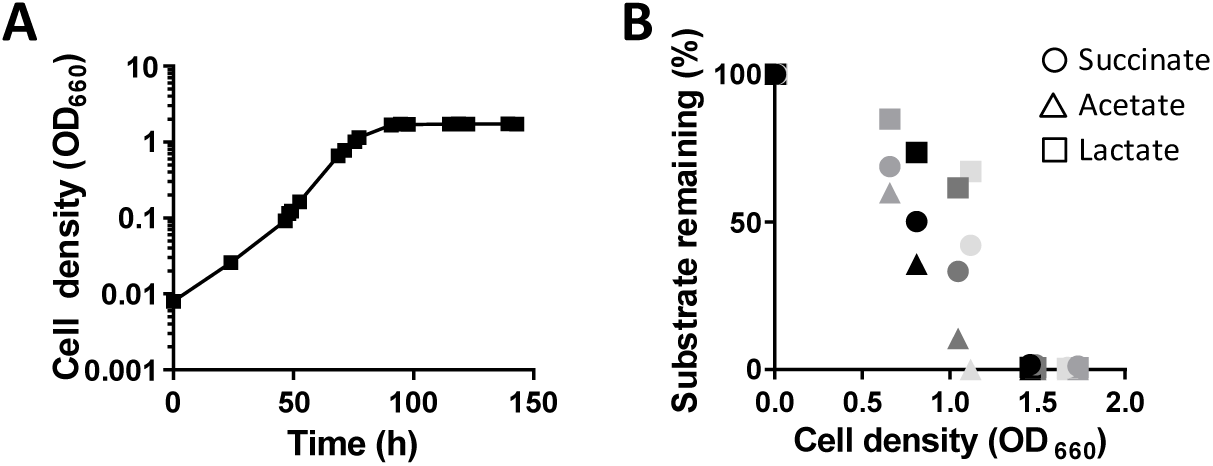
*R. palustris* Nx simultaneously utilizes succinate, acetate, and lactate when provided a mix of the three substrates. (**A**) Representative growth curve of *R. palustris* Nx grow in MDC with 5 mM each succinate, acetate, and lactate. Similar trends were observed for three other biological replicates (**B**) Amount (%) of succinate (circles), acetate (triangles), and lactate (squares), remaining in culture supernatants at the indicated cell densities. Each of the four shades of gray indicates an independent biological replicate.

To ascertain if lactate consumption was stimulated by both, one, or either succinate or acetate, we also examined growth and carbon utilization in cultures pairing lactate with either acetate or succinate. Cultures reached stationary phase with each of these mixtures within 150-200 h (**Fig. 2A**). For comparison, no growth was detected during the same time period when 10 mM lactate was provided as the sole carbon source (**Fig. 2A**). The percentages of carbon consumed (**Fig. 2B**) and final cell densities per mol carbon consumed (**Fig. 2C**) were comparable between cultures provided one, two, or three substrates, indicating that all substrates were being assimilated into biomass in these cultures.

**Fig. 2.**
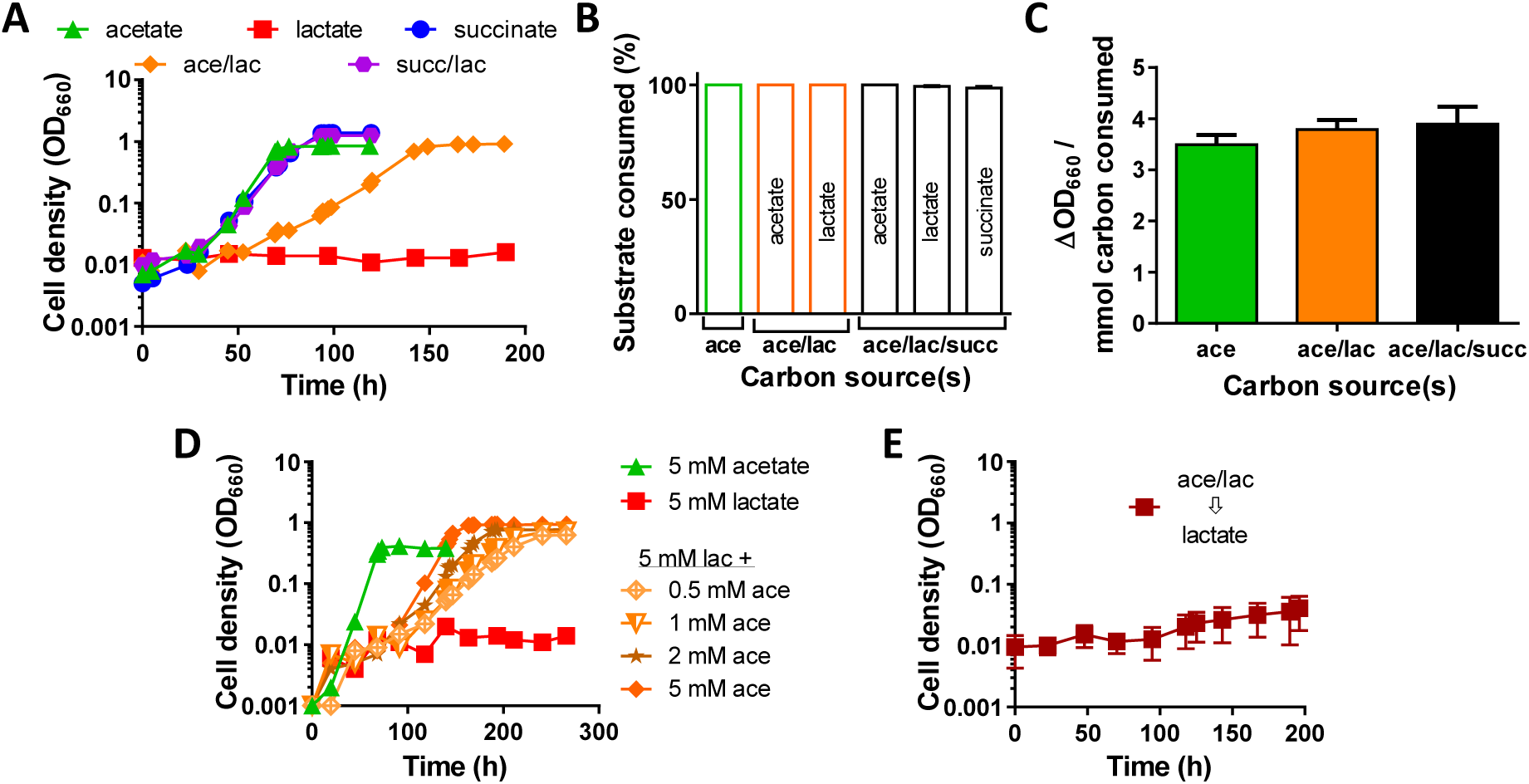
Both succinate and acetate individually stimulate expedited co-utilization of lactate by *R. palustris* Nx. (**A**) Representative growth curves of *R. palustris* Nx in MDC with 10 mM individual substrates (acetate [ace], lactate [lac], or succinate [succ]) or 5 mM each of paired substrates (ace/lac, succ/lac). (**B, C**) Amount (%) of each substrate consumed at stationary phase (**B**) and OD yields (**C**) of cultures provided 10 mM acetate, 5 mM each acetate and lactate, or 5 mM each succinate, acetate, and lactate. Error bars, SD; n≥3. (**D**) Representative growth curves of *R. palustris* Nx in MDC with 5 mM acetate alone, 5 mM lactate alone, or 5 mM lactate supplemented with the indicated concentrations of acetate. (**A, D**) Similar trends were observed for three other biological replicates for each condition. (**E**) Growth curve of *R. palustris* Nx that was subcultured from stationary-phase ace/lac cultures into MDC with 10 mM lactate alone. Error bars, SD; n=4.

To assess the sensitivity of *R. palustris* lactate utilization to the presence of co-substrate, we next examined growth in cultures containing a 5mM lactate with varying amounts of acetate (0.5mM – 5mM). Although the growth rates varied slightly with acetate concentration, stimulation of lactate utilization occurred with all acetate concentrations tested (**Fig. 2D**). Thus, lactate utilization can be stimulated by a range of co-substrate concentrations and different lactate:co-substrate ratios. Notably, cells from cultures grown with acetate plus lactate, which presumably contained the necessary enzymes for lactate catabolism, failed to grow when transferred to fresh medium with lactate as the sole carbon source (**Fig. 2E**), suggesting that co-utilization itself was necessary for expedited lactate utilization and that co-substrates are inadequate to prime cell physiology for growth on lactate as the sole carbon source. Based on these data, we conclude that utilization of acetate or succinate is sufficient to stimulate the expedited and simultaneous co-utilization of lactate by *R. palustris*.

### Mixed-substrate utilization stimulates lactate consumption in diverse *R. palustris* strains

The above experiments examined lactate utilization under conditions that mimicked coculture conditions, wherein lactate co-utilization was first observed, in the following two regards. First, we used the engineered ‘Nx’ strain of *R. palustris*, which harbors a mutation in *nifA* resulting in constitutive N_2_ fixation, deletion of *hupS* to prevent H_2_ oxidation, and deletion of *hfsE* to prevent cell aggregation (21, 22). Second, the cultures were grown in a minimal medium (MDC) with N_2_ as the sole nitrogen source (21). To assess if the engineered mutations and/or N_2_-fixing conditions contributed to the co-utilization of lactate with acetate and succinate, we examined carbon utilization in CGA009, the wild-type parent strain of *R. palustris* Nx, grown with the acetate, succinate, and lactate mixture or with lactate alone in either MDC or in an NH_4_^+^-containing minimal medium, PM. The presence of NH_4_^+^ in PM represses N_2_-fixation in CGA009 (23). Lactate utilization patterns were similar to those in *R. palustris* Nx, regardless of the media: CGA009 consumed all three compounds when provided as a mixture within 150 h and failed to grow with lactate alone in the same time frame (**Fig. 3**). Thus, the observed lactate consumption patterns were not due to either the engineered mutations in the Nx strain or the N_2_-fixing conditions.

**Fig. 3.**
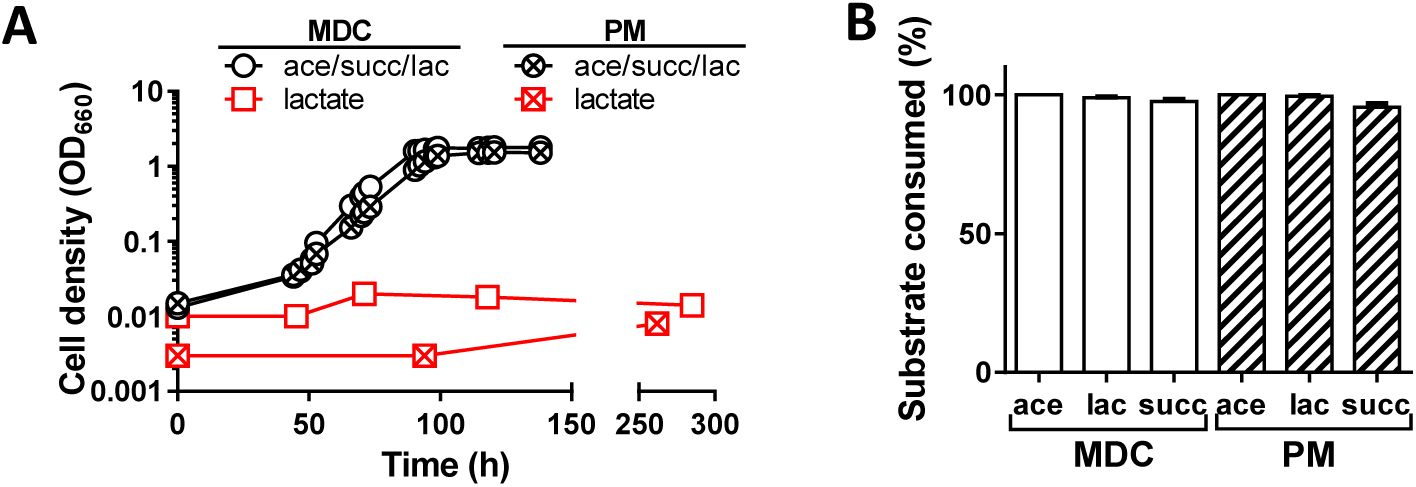
Stimulation of lactate consumption via co-utilization of acetate and succinate also occurs in wild-type *R. palustris* CGA009 and is independent of N_2_ fixation. (**A**) Representative growth curves of *R. palustris* CGA009 in MDC or PM with either 5 mM each succinate, acetate, and lactate (ace/succ/lac) or 10 mM lactate alone. Similar trends were observed for three other biological replicates in each condition. (**B**) Amount (%) of each substrate consumed at stationary phase in cultures of *R. palustris* CGA009 in MDC or PM with 5 mM each succinate, acetate, and lactate. Error bars, SD; n=4.

We also investigated if acetate and succinate stimulated lactate utilization in environmental *R. palustris* strains. Environmental isolates of *R. palustris* have large genetic differences and exhibit unique metabolic characteristics that are thought to aid in nutrient acquisition, anaerobic fermentation, and/or light-harvesting (24-26). Thus, it was conceivable that other *R. palustris* strains behave differently with regard to lactate utilization, either readily using lactate as a sole carbon source or failing to use lactate even in the presence of additional organic acids. However, these potential alternatives were refuted for two environmental isolates, namely, BisB5 and DX-1. When BisB5 and DX-1 were grown with acetate, succinate, and lactate in PM (**Fig. 4A**), all three compounds were consumed within 120 hours (**Fig. 4B**). In contrast, little or no growth was observed with lactate as the sole carbon source within the same time frame (**Fig. 4A**). These results indicate that stimulation of lactate catabolism via mixed-substrate utilization is conserved among diverse *R. palustris* strains.

**Fig. 4.**
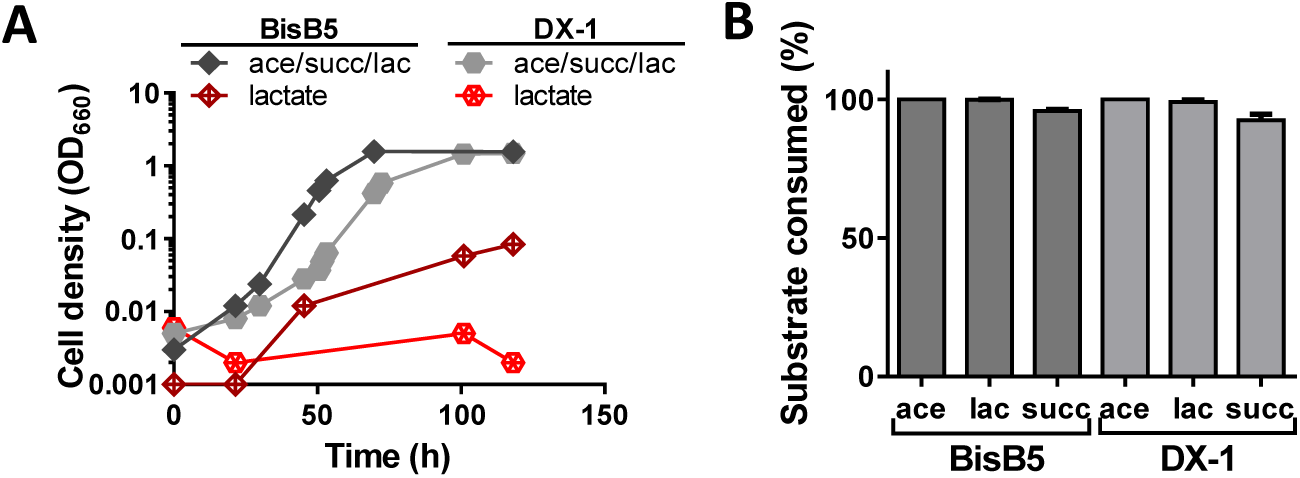
Stimulation of lactate consumption via co-utilization of acetate and succinate also occurs in environmental *R. palustris* isolates. (**A**) Representative growth curves for *R. palustris* strains BisB5 or DX-1 in PM with either 5 mM each succinate, acetate, and lactate or 10 mM lactate alone. Similar trends were observed for three other biological replicates for each strain in each condition.(**B**) Amount (%) of each substrate consumed at stationary phase in cultures of *R. palustris* BisB5 or DX-1 in PM with 5 mM each succinate, acetate, and lactate. Error bars, SD; n=4.

### *R. palustris* Nx lactate utilization is stimulated by diverse carbon co-substrates

The carbon substrates available to *R. palustris* in natural environments are presumably more diverse than *E. coli* fermentation products. Therefore, we investigated if lactate utilization by *R. palustris* was stimulated by co-consumption of carbon substrates other than acetate and succinate. Specifically, we grew *R. palustris* with malate, butyrate, and glycerol, as either the sole carbon source or paired with lactate. Glucose was not tested because *R. palustris* cannot consume sugars (27). Similar to the results with acetate and succinate, lactate was utilized simultaneously with malate, butyrate, and glycerol (**Fig. 5A-C**). These data demonstrate that lactate utilization can be stimulated by diverse co-substrates.

**Fig. 5.**
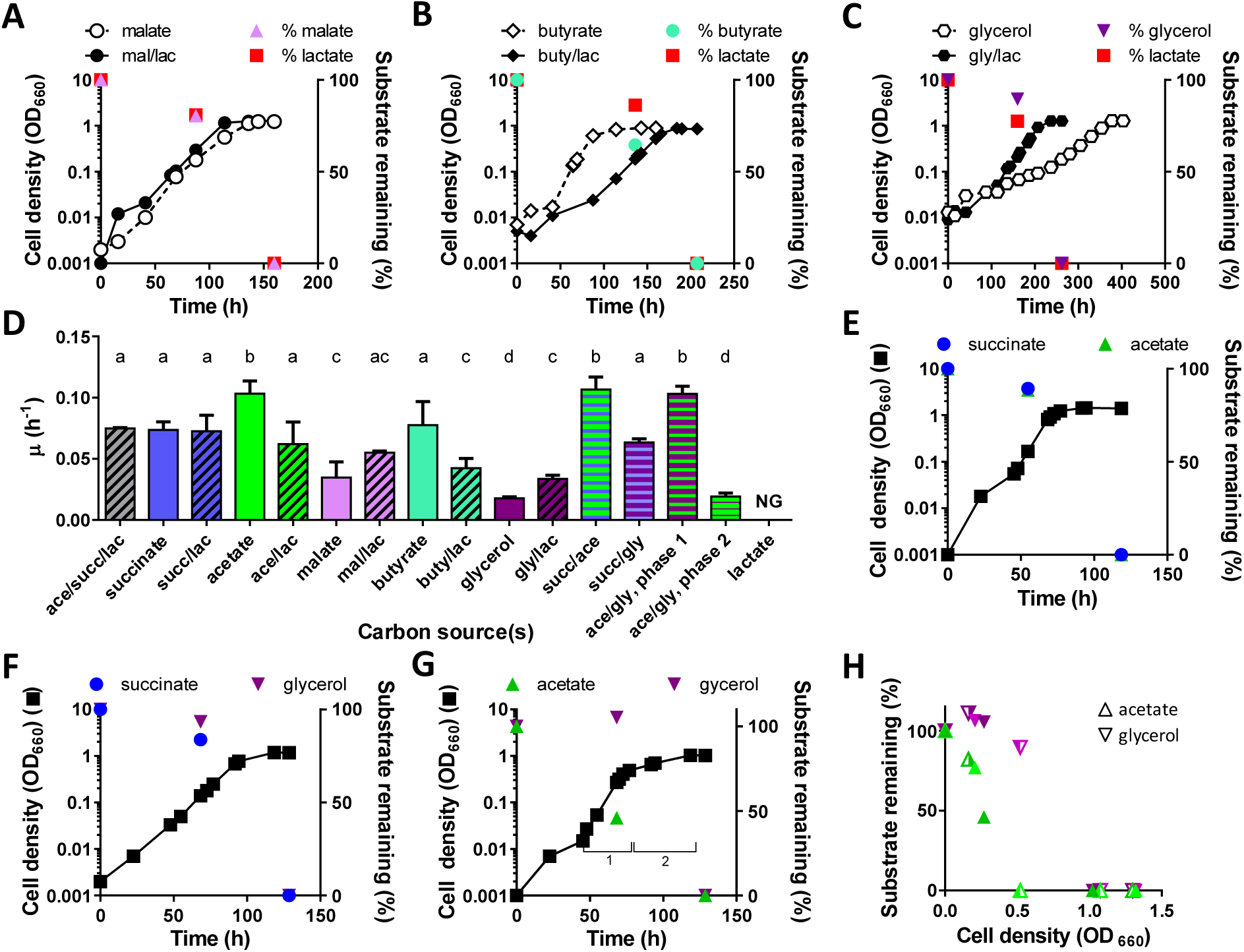
*R. palustris* co-utilizes many, but not all, carbon substrate pairs. (**A-C**) Representative growth curves and amount (%) of substrates remaining at early-log phase for *R. palustris* Nx in MDC with 5 mM each of the following paired substrates: malate + lactate (mal/lac) (**A**), butyrate + lactate (buty/lac) (**B**), and glycerol + lactate (gly/lac) (**C**). Similar trends were observed for two or more additional biological replicates for each condition. (**D**) Specific growth rates of *R. palustris* Nx in MDC with indicated carbon substrates. NG, no growth. Error bars, SD; n≥3. Different letters indicate statistically significant differences between groups (P < 0.05; one-way ANOVA with Tukey’s multiple-comparison test). (**E-G**) Representative growth curves and amount (%) of substrates remaining at log phase and stationary phase for *R. palustris* Nx in MDC with 5 mM each of the following paired substrates: succinate + acetate (succ/ace) (**E**), succinate + glycerol (succ/gly) (**F**), and acetate + glycerol (ace/gly) (**G**). Similar trends were observed for two or more additional biological replicates. (**G**) The numbered brackets indicate the two exponential growth phases (see **D**). (**H**) Amount (%) of acetate and glycerol remaining in supernatants of ace/gly cultures at indicated cell densities. Each of the four shades of green (acetate) or purple (glycerol) indicates an independent biological replicate.

### Differential effects of mixed-substrate utilization on *R. palustris* Nx growth

In working with different substrate mixtures containing lactate, we noticed that co-utilization sometimes resulted in different growth rates compared to the growth rates on single carbon sources. Specific growth rates (µ) during co-utilization could be categorized in comparison to the growth rates on the constituent substrates alone, as follows: (i) the mixed-substrate µ was faster than that on either substrate alone (i.e., enhanced µ); (ii) the mixed-substrate µ approximated that when grown on the individual substrate allowing the fastest growth (i.e., equivalent µ); or (iii) the mixed-substrate µ was between the µ’s on the individual substrates (intermediate µ). We considered cultures with lactate as the sole carbon source to have a growth rate of 0 h^-1^, as no growth was observed in these cultures within experimental time frames (≤ 300 h). We observed enhanced µ in cultures pairing lactate with glycerol, equivalent µ in cultures pairing lactate with succinate or malate, and intermediate µ in cultures pairing lactate with acetate or butyrate (**Fig. 5D**). Akin to growth patterns in other species (13), there was no evident correlation between the effect of mixed-substrate utilization on growth rate and either the metabolic entry point of the co-substrate or the growth rate on the co-substrate alone.

To investigate if the changes in mixed-substrate growth rates were contingent on the co-substrate rather than on lactate itself, we examined growth of *R. palustris* with three substrate pairs that did not contain lactate: succinate with acetate, succinate with glycerol, and acetate with glycerol. When acetate was paired with succinate, the compounds were utilized simultaneously and the growth rate matched that of cultures with acetate alone (equivalent µ) (**Fig. 5D,E**). Similar results were seen in cultures containing succinate paired with glycerol, with growth rates approximating those of succinate, the ‘preferred’ carbon source (equivalent µ) (**Fig. 5D,F**). However, pairing acetate with glycerol resulted in two distinct exponential growth phases, with the first and second phases having growth rates that approximated those with acetate alone and glycerol alone, respectively (**Fig. 5D,G**). This pattern suggested that acetate and glycerol were being consumed sequentially, rather than simultaneously. HPLC results confirmed that acetate consumption occurred during the first exponential phase whereas glycerol consumption did not occur until acetate had been depleted from the medium (**Fig. 5G,H**). From these data, we conclude that, whereas *R. palustris* can simultaneously consume a wide range of substrates when provided in mixtures of two and three, acetate and glycerol are consumed sequentially by *R. palustris*.

## DISCUSSION

Here we revealed that lactate can be readily catabolized by *R. palustris* in the presence of various other organic acids and glycerol (**Figs. 2 and 5**), despite that lactate did not support growth as the sole carbon source in the same time frames. It is tempting to speculate how co-utilization expedites lactate consumption. The fact that we observed a similar induction effect with diverse substrates that enter central metabolism at different points of both glycolysis/gluconeogenesis and the TCA cycle makes it difficult to predict the underlying mechanism(s). However, we believe several mechanisms can be excluded. First, there are instances where co-substrates enable anaerobic growth by acting as alternative electron acceptors and thereby contributing to cellular redox balance (28-30). The contribution of co-substrates to electron balance during lactate co-utilization is unlikely because: (i) the same pattern of lactate utilization was observed in two conditions that differentially allow N_2_ fixation (**Fig. 3**), a process known to satisfy electron balance in *R. palustris* (16); and (ii) lactate utilization was stimulated equivalently by carbon substrates that were more oxidized or less oxidized than lactate (**Table 1**). Second, co-transport is unlikely to be responsible, as co-consumption was not strictly dependent on lactate:co-substrate stoichiometry (**Fig. 2D**) and induction occurred with diverse co-substrates that presumably do not all utilize the same transporter (**Figs. 2 and 5**). Finally, in some instances co-substrates can have an “auxiliary” effect by providing energy during the catabolism of energy-deficit substrates (1, 31). The need for supplemental energy generation is unlikely in the case presented herein, given that *R. palustris* was grown under phototrophic conditions where energy is derived from light. Although outside of the scope of this study, we hope that future work identifies the mechanism(s) by which mixed-substrate utilization expedites lactate consumption by *R. palustris*.

**Table 1.** Oxidation states of tested *R. palustris* growth substrates

| Substrate | Formula | Oxidation state<sup>a</sup> |
| --- | --- | --- |
| Malate | C<sub>4</sub>H<sub>6</sub>O<sub>5</sub> | +1 |
| Succinate | C<sub>4</sub>H<sub>6</sub>O<sub>4</sub> | +0.5 |
| Lactate | C<sub>3</sub>H<sub>6</sub>O<sub>3</sub> | 0 |
| Acetate | C<sub>2</sub>H<sub>3</sub>O<sub>2</sub> | 0 |
| Glycerol | C<sub>3</sub>H<sub>8</sub>O<sub>3</sub> | -0.67 |
| Butyrate | C<sub>4</sub>H<sub>8</sub>O<sub>2</sub> | -1 |

<sup>a</sup>Oxidation states were calculated as previously described (19)

This study was initiated to investigate the potential co-utilization of lactate with other carbon substrates. However, our results also revealed acetate inhibition of glycerol catabolism in *R. palustris* (**Fig. 5G, H**). *R. palustris* is well known for its strict control of nitrogen utilization, wherein the presence of ammonium strictly inhibits expression of the nitrogenase enzyme that catalyzes N_2_ fixation (23, 32). However, we are unaware of any report of CCR in this species. Notably, there was no evident lag phase between the two exponential growth phases in *R. palustris* cultures containing acetate paired with glycerol (**Fig. 5G**). The lack of an intervening lag phase suggests that acetate-mediated inhibition of glycerol assimilation in *R. palustris* may be occurring at the level of protein activity (e.g., transport or catabolic enzyme activity), rather than the level of protein expression (2, 4, 33). *R. palustris* CGA009 has more than 400 genes predicted to be involved in regulation and signal transduction (27). Among these are genes encoding Crp- and Hpr-like proteins (27, 34). Crp and Hpr homologues regulate diverse biological functions that include CCR in certain species (34, 35). As such, the Hpr- and Crp-like proteins seem logical initial targets for mutagenesis in the endeavor to characterize catabolite control mechanisms in *R. palustris*. Identifying the transporters used for different carbon substrates in *R. palustris* will likely also be important for elucidating such mechanisms. As *R. palustris* encodes more than 300 different transport systems (27), and results from a large-scale study of ABC transporter proteins indicate that sequence-based homology is unreliable for predicting ligand specificity (36), this will not be a trivial task.

Although simultaneous utilization of carbon substrates is most commonly described under nutrient-limited conditions (2-4), examples are accumulating, including for *R. palustris* as shown here, wherein bacteria simultaneously utilize substrates even at high concentrations (1, 13). *R. palustris* simultaneously consumed seven of the eight substrate pairs tested in this study, and published data suggest that this behavior may extend beyond organic acids and glycerol. For example, data from a recent study indicated that *R. palustris* simultaneously utilizes acetate and various aromatic compounds when grown in corn stover hydrolysate (20), though it was not determined which compounds were being assimilated into biomass. The same study reported simultaneous biological transformation of several aromatic compounds that are not readily utilized as sole carbon sources (15, 20), perhaps indicating that mixed-substrate utilization influences the aromatic utilization spectrum of *R. palustris* as well. It is possible that assessment of bacterial nutritional repertoires using single substrates underestimates the catabolic capabilities of some bacteria. From an ecological perspective, it would not necessarily be surprising if *R. palustris* co-utilizes a large range of carbon sources. Such a strategy could allow *R. palustris* to take full advantage of the diverse carbon sources it encounters within the numerous environments it inhabits (27, 37). It has been proposed that carbon source preference reflects the likelihood of encountering various substrates in the environment (38). Thus, to speculate further, the disparity between lactate utilization in the presence and absence of a co-substrate could indicate that lactate is rarely encountered as the sole carbon source in natural environments. In this case, the inability to readily use lactate as the sole carbon source would not be of consequence to *R. palustris*. Finally, beyond these potential ecological implications, substrate co-utilization, particularly at high substrate concentrations, is preferable for industrial and commercial applications (8, 39). Specifically, such behavior is crucial for developing bioprocesses that utilize cheap, renewable waste materials, such as industrial effluents, lignocellulosic biomass, and food waste, as feedstocks for the production of biofuels and value-added products. We believe the proclivity to co-utilize carbon substrates enhances the potential biotechnological value of *R. palustris*.

## MATERIALS AND METHODS

### Chemicals, strains, and growth conditions

The *R. palustris* strains used in this study are listed in Table 2. *R. palustris* was routinely cultivated on defined mineral (PM) (40) agar supplemented with 10 mM succinate. All cultures were grown in 27-mL anaerobic test tubes containing 10 mL of either defined M9-derived coculture medium (MDC) (21) or PM medium. MDC or PM was made anaerobic by bubbling with 100% N_2_ or Ar, respectively, and then sealing with rubber stoppers and aluminum crimps prior to autoclaving.

**Table 2.** *R. palustris* strains used in this study.

| Strain | Description or Sequence (5'-3'); Paper designation | Reference |
| --- | --- | --- |
| CGA009 | Wild-type strain; spontaneous Cm<sup>R</sup> derivative of CGA001 | (40) |
| CGA4005 | CGA009 *nifA* * $\Delta$ *hupS* $\Delta$ *rpa2750*; Nx | (21) |
| BisB5 | Environmental isolate | (24) |
| DX-1 | Environmental isolate | (42) |

For starter cultures, single colonies were used to inoculate MDC with limiting (3 mM) acetate. For experimental cultures, 100 µL aliquots of replicate stationary-phase starter cultures were used to inoculate MDC or PM supplemented with either 10 mM of a single carbon substrate or 5 mM each of multiple carbon substrates, unless indicated otherwise in figure legends. Carbon sources were added to desired final concentrations from 1M stock solutions of glycerol and sodium salts of L-lactate, acetate, succinate, L-malate, and butyrate. All cultures were incubated horizontally at 30°C under a 43 W A19 halogen bulb (750 lumens) with shaking at 150 rpm. At least three independent biological replicates were performed for each culture condition.

### Analytical procedures

*R. palustris* growth was monitored via optical density at 660 nm (OD_660_) using a Genesys 20 spectrophotometer (Thermo-Fisher, Waltham, MA, USA). Growth readings were measured in culture tubes without sampling. Specific growth rates were calculated using OD_660_ values between 0.1—1.0 where cell density and OD_660_ are linearly correlated. Final cell densities were measured in cuvettes with samples diluted as needed to achieve an OD_660_ within the linear range. Organic acids and glycerol were quantified using a Shimadzu high-performance liquid chromatograph, as previously described (41).

## ACKNOWLEDGEMENTS

The authors thank Julia van Kessel, Ankur Dalia, and members of the McKinlay Lab for discussions. This work was supported in part by the U.S. Department of Energy, Office of Science, Office of Biological and Environmental Research, under award DE-SC0008131, the U.S. Army Research Office Grant # W911NF-14-1-0411, and the National Science Foundation CAREER award # MCB-1749489.

## Notes

Conflict of interest: The authors declare no conflict of interest.

